# Biotic interactions limit the geographic range of an annual plant: herbivory and phenology mediate fitness beyond a range margin

**DOI:** 10.1101/300590

**Authors:** John W. Benning, Vincent M. Eckhart, Monica A. Geber, David A. Moeller

## Abstract

Species’ range limits offer powerful opportunities to study environmental factors regulating distributions and probe the limits of adaptation. However, we rarely know what aspects of the environment are actually constraining range expansion, much less which traits are mediating the organisms’ response to these environmental gradients. Though most studies focus on climatic limits to species’ distributions, biotic interactions may be just as important. We used field experiments and simulations to estimate contributions of mammal herbivory to a range boundary in the annual plant *Clarkia xantiana* ssp. *xantiana.* A steep gradient of increasing probability of herbivory occurs across the boundary, and herbivory drives several-fold declines in lifetime fitness at and beyond the boundary. By including in our analyses data from a sister taxon with more rapid phenology, we show that delayed phenology drives *C. xantiana* ssp. *xantiana’s* susceptibility to herbivory and low fitness beyond its border.

## Introduction

Understanding the causes of species’ geographic range limits is a fundamental problem in ecology and evolution. For the vast majority of species, however, we still cannot answer why an organism occurs on one side of its range boundary and not the other (Gaston 2009). Pinpointing the underlying environmental drivers and demographic and genetic mechanisms restricting species distributions is of utmost importance for understanding species’ responses to global change (Alexander *et al.* 2015; Ettinger & HilleRisLambers 2017), the spread of invasive species (Colautti *et al.* 2010), and the limits to natural selection (Antonovics 1976; Kawecki 2008).

Some species have simply not had time to colonize environmentally suitable areas (dispersal lag; Svenning *et al.* 2008; Alexander *et al.* 2017), and in other cases, abrupt dispersal barriers can prevent range expansion (Chardon *et al.* 2015; Weir *et al.* 2015). However, most species’ borders occur along seemingly gradual environmental gradients (Kirkpatrick & Barton 1997; Sexton *et al*. 2009) and are linked to underlying variation in the environment across the landscape and corresponding variation in adaptation. Species may be restricted to their current distribution simply because they are maladapted to the environment beyond their range boundary.

Several theoretical models address the apparent “failure” of natural selection to result in adaptation to novel environments outside a species’ range (e.g., Kirkpatrick & Barton 1997; Case & Taper 2000; Polechová & Barton 2015). Population dynamics in these models are based upon the difference between a population’s realized value of some important trait, and the optimal trait value dictated by the environment; this difference determines the degree of population maladaptation and population growth (Kirkpatrick & Barton 1997).

A key factor in these models of range limits is the steepness of the environmental gradient along which populations must adapt. As gradients become steeper, adaptation to areas outside the current range becomes less likely due to high levels of maladaptation in colonists dispersing from the range edge; with shallow clines, adaptation and expansion of the range limit can proceed (Kirkpatrick & Barton 1997; Polechová & Barton 2015). Most models assume linear gradients in environmental variables, but non-linear gradients can be especially important in generating distributional limits due to rapid change in optimal phenotype across space (Case & Taper 2000; Polechová & Barton 2015).

Given the central role of environmental gradients in structuring species’ distributions, identifying important gradients is usually a first goal of range limit studies, with climatic variables being likely candidates. While climatic niche limits often do explain species’ distributions (Lee-Yaw *et al*. 2016), it is increasingly recognized that biotic interactions can contribute to large scale distributional limits (Case *et al.* 2005; Araújo & Luoto 2007; Hargreaves *et al.* 2014; Louthan *et al.* 2015). Theoretical and empirical work has shown that competition (Case & Taper 2000; Ettinger & HilleRisLambers 2017), predation (Bruelheide & Scheidel 1999; deRivera *et al.* 2005), parasitism (Hochberg & Ives 1999; Briers 2003), and mutualism (Stanton-Geddes & Anderson 2011; Moeller *et al.* 2012; Afkhami *et al.* 2014; Lankau & Keymer 2016) can mediate species’ range limits.

Though correlative approaches such as species distribution models lend first insights into potential drivers of range limits, transplant experiments including sites outside the range limit are the only way to test range-boundary hypotheses directly (Hargreaves *et al.* 2014). Transplant experiments paired with field measurements of potentially important traits may reveal trait-environment relationships that underlie geographic variation in performance (Hoffmann & Blows 1994; Angert *et al.* 2008; Sexton *et al.* 2009; Hargreaves *et al.* 2014).

Investigating ecological causes of a species’ distributional limit thus has three main components: characterizing environmental gradients, linking gradients to individual and population fitness, and determining the trait(s) mediating fitness responses. Studies rarely tackle these three points in concert (but see Angert *et al*. 2008), especially in regard to biotic interactions. Here we investigate the role of an antagonistic interaction, mammal herbivory, in limiting the range of an annual plant, *Clarkia xantiana* ssp. *xantiana.* With two years of stem-translocation experiments, we determined the shape of the gradient in fatal herbivory across a major range boundary. Reanalyzing data from a previous reciprocal transplant experiment across the same boundary (Geber & Eckhart 2005), we calculated the magnitude of mammalian herbivory over *xantiana’*s full lifespan, and used those estimates in new simulations of herbivory’s effects on fitness that estimated how fitness would change if herbivory were less intense or absent. Finally, we tested the hypothesis that susceptibility to herbivory depends on a specific plant trait, phenology, quantifying the relationship of phenology to herbivory risk across the range boundary.

## Material and methods

### Study System

*Clarkia xantiana* is comprised of two subspecies, *C. x.* ssp. *xantiana* A. Gray and *C. x.* ssp. *parviflora* (Eastw.) Harlan Lewis and P.H. Raven (hereafter, *xantiana* and *parviflora).* Both taxa are annuals endemic to the foothills of the southern Sierra Nevada and Transverse Ranges of California (Moore & Lewis 1965; Eckhart & Geber 1999). Their combined range spans a complex west-to-east environmental gradient with *xantiana* found in the wetter western region in oak woodlands, and *parviflora* found in the eastern region in arid scrub and pinyon-juniper woodland (Fig. 1; (Eckhart & Geber 1999)).

**Figure 1.**
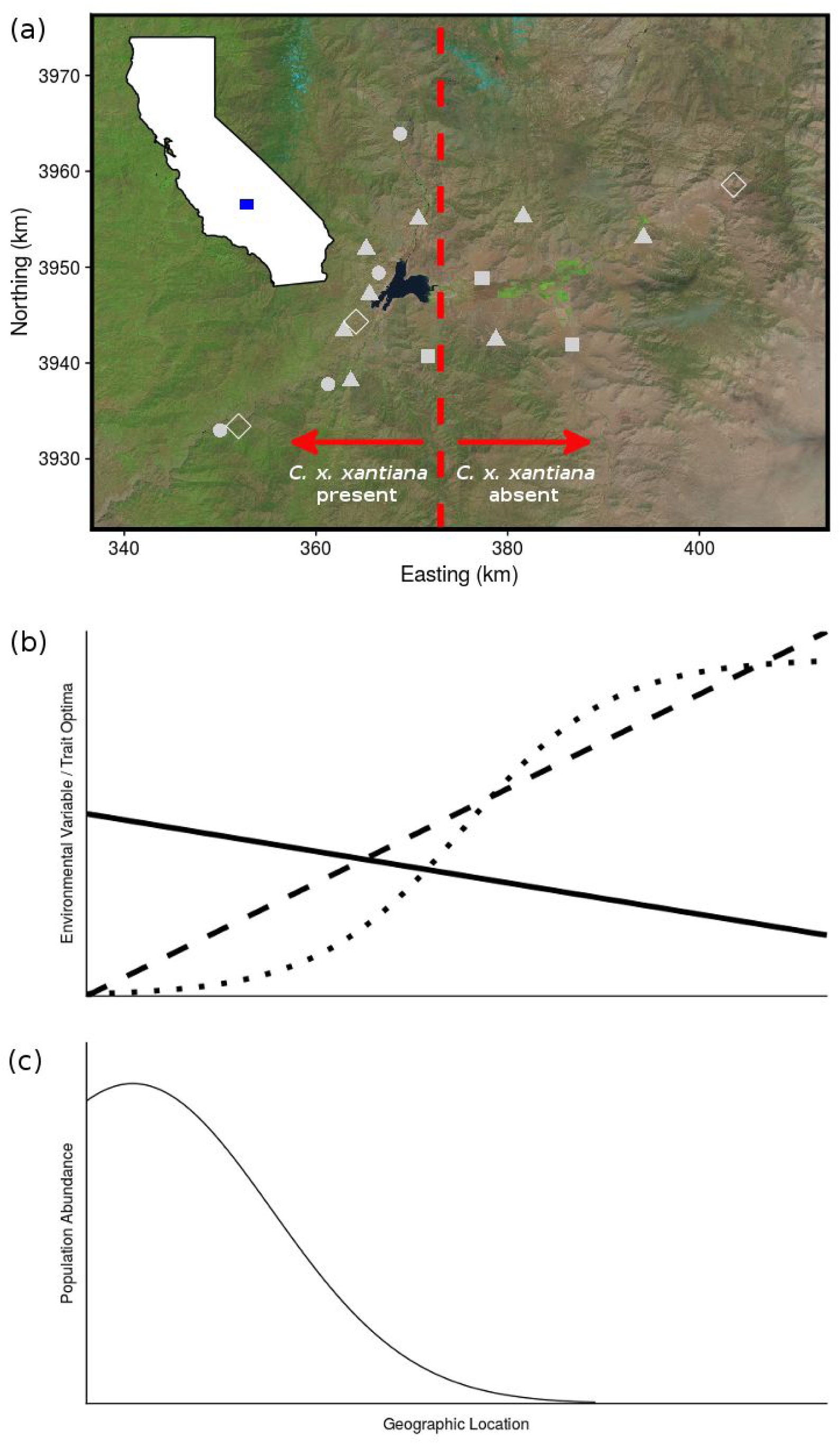
Study area and conceptual overview. (a) shows satellite photo of study area in the Southern Sierra Nevada foothills, with inset highlighting location within California. Red line marks *C. x. xantiana*’s eastern range limit. Also shown are locations of 2015-2016 stem translocation sites (filled symbols; circles mark sites used in 2015, triangles mark sites used in 2016, squares mark sites used in both years) and 1997-1999 reciprocal transplant sites (open diamonds). Panels (b) and (c) illustrate how gradients in environmental variables, and thus clines in trait optima for organisms, can regulate population abundance across a species’ range. Panel (b) illustrates how multiple environmental gradients shifting across a species’ distribution can vary in their shape and slope, exhibiting linear increases or decreases, or regions with abrupt changes in gradient steepness. The steeper the cline, the less likely adaptation will enable peripheral populations to match trait optima and colonize areas further outside the current range limit; nonlinear clines can be especially important in setting range limits. Panel (c) plots population abundance across space as controlled by these environmental gradients via the degree of mismatch between populations’ trait means and the local trait optima; a range limit forms where abundance drops to zero. (b) and (c) modified from (<u>Kirkpatrick and Barton 1997</u>).

The two taxa are in secondary contact (in a narrow zone of sympatry) after diverging ca. 65,000 years ago (Pettengill & Moeller 2012a, b), and have differentiated most strongly in mating system and phenology (Eckhart & Geber 1999). Both are self-compatible, but *parviflora* is predominantly selfing, while *xantiana* is predominantly outcrossing (Moeller 2006). *Parviflora* also completes its life cycle more quickly than *xantiana*, and this phenological differentiation contributes to the near complete reproductive isolation between the subspecies (Briscoe Runquist *et al.* 2014). A reciprocal transplant experiment showed each subspecies to be strongly locally adapted to its own home range (Geber & Eckhart 2005). For *xantiana*, a west-to-east gradient of increasing aridity (Eckhart *et al.* 2010) and declining pollinator abundance and diversity (Moeller 2005, 2006; Moeller *et al.* 2012) contribute to reducing *xantiana* fitness below population-sustaining levels beyond its eastern range edge (Geber & Eckhart 2005). In the transplant study, there was preliminary evidence that small mammal herbivory influenced *xantiana*’s performance beyond its range edge.

### Quantifying patterns of herbivory across and beyond the range

To identify fine scale spatial and temporal variation in plant-herbivore interactions, we performed a stem-translocation experiment across two years at 15 sites inside and outside *xantiana’s* range. Clipping living adult stems from natural populations to create experimental arrays, we quantified herbivory while controlling for genotype, plant size, and phenology. Experiments were conducted in or near to natural *xantiana* and *parviflora* populations. In 2015, we quantified broad-scale variation in herbivory across most of the west-east extent of *xantiana’s* range and beyond the range limit. We sourced *xantiana* stems from the center of the range, and within 6 km of the eastern edge. We placed stems at seven sites (two at range center, three at range edge, two that were 5 and 14 km beyond the eastern range limit; Fig. 1A). At each site we installed two transects of 24 stems, alternating central and edge genotypes, with stems placed 1 m apart. Plant stems were maintained in 13 cm florist picks filled with water and secured with an attached metal rod sunk into the ground (Fig. S1 A). Plants maintained in this way continue to open new flowers and set fruits after pollination (JW Benning, pers. obs.). To explore temporal variation in herbivory, we installed four temporal replicates of stems in May and June (approximately once per week from 24 May - 19 June) at each site. For each temporal replicate we scored stems for fatal herbivory (having no buds, flowers, or fruits remaining, usually because most of the stem was completely removed) five days after installation (Fig. S1D). At the five sites within *xantiana’s* range, we also followed naturally occurring plants near experimental arrays to determine whether geographic patterns of herbivory on experimental plants mimicked that on natural plants (Appendix 1).

Our 2015 experiment showed that herbivory was low in the range center and much stronger at the range edge and beyond; however, the coarse geographic scale covered did not allow for a fine-scale characterization of the environmental gradient at the range limit. In 2016, we established experimental arrays in six sites near to or at the range limit, and five sites outside the range limit (Fig. 1A). As the 2015 experiment showed no effect of population source (central vs edge genotypes), plants used in 2016 were a mixture of genotypes from across the range. At each site we installed three transects of 10 stems placed 1 m apart. In an attempt to further mimic natural plant conditions, we placed 2016 stems in 50 mL conical tubes sunk completely into the ground (Fig. S1B). We installed three temporal replicates of stems at each site and scored herbivory five days after installation. In 2016, wildfires destroyed the third round of experimental stems at three sites.

We used logistic regression to test the effects of easting (i.e., longitude), time (temporal replicate), and, in 2015, genotype source (central vs. edge), and all interactions, on the probability of herbivory. For both years, transect was included as a term nested within census date and easting position. Models were constructed using the glm function in R (R Core Team 2017), with binomial error distribution and logit link. We used BIC (Bayesian Information Criterion) scores to compare models with linear, linear plus quadratic, and linear plus quadratic plus cubic easting terms. We tested the significance of each term using Type II ANOVAs with likelihood ratio tests (car package; Fox & Weisberg 2011) and calculated model R^2^ (Nagelkerke’s pseudo R^2^) using the sjstats package (Lüdecke 2018). Conditional predicted probabilities were calculated and plotted with the visreg (Breheny & Burchett 2017) and ggplot2 (Wickham 2009) packages in R.

### Quantifying the effects of herbivory on population fitness

We used data from a two-year reciprocal transplant experiment to ask how herbivory affects population fitness and the likelihood of population persistence across and beyond the range limit of *xantiana.* We compared our results for *xantiana* to those of *parviflora* (its sister taxon beyond *xantiana’*s eastern range limit) as a means of identifying how trait differences between the two taxa may result in differing performance and susceptibility to herbivory.

### Reciprocal transplant

In 1997-1999, we conducted a reciprocal transplant experiment to examine variation in phenotypic traits and lifetime fitness of both subspecies planted within and outside their respective ranges. In each year of this experiment, we planted 6 populations of *xantiana* and 12 populations of *parviflora* at one site within *xantiana’*s range center (but outside *parviflora’*s western range limit; Center), one site at *xantiana*’s range edge where it overlaps narrowly with *parviflora*’s range (Edge), and one site beyond the eastern *xantiana* range limit (but within *parviflora*’s distribution; Beyond-Edge; Fig. 1A). We planted seeds into 8,488 planting positions (eight seeds per position) in October and scored germination and survival monthly from January through July. The two years of the experiment differed markedly in precipitation, and this led to strong differences in lifetime fitness estimates between years; hereafter, we refer to the two years of the experiment as “wet” (1997-1998) and “dry” (1998-1999). Full experimental details can be found elsewhere (Eckhart *et al.* 2004; Geber & Eckhart 2005).

#### Simulation of fitness in the absence of herbivory

We first simulated a scenario where there was no fatal mammalian herbivory during the two-year field experiment. In essence, we took the original experiment’s data set and, for each plant that suffered fatal herbivory, estimated how many seeds it would have produced had it not been eaten. We used predictive models built with field data to produce lifetime fitness estimates for eaten plants that reflected all other environmental aspects of the sites, while “removing” herbivory. Predictive models were evaluated using R^2^ statistics and diagnostic plots of predicted versus observed values (see Appendix S1 for details on model construction).

After calculating predicted fitness values for eaten plants, we examined the extent to which average lifetime fitness would change at each site if there was no fatal mammalian herbivory. We estimated average lifetime fitness through female function (seeds produced per planted seed) for each subspecies at each site in both years. We used linear mixed models of lifetime fitness with site, year, and subspecies as fixed factors, and block (nested within site and year) and population (nested within subspecies) as random factors (as in Geber and Eckhart 2005). Comparison of the least-square means from models based on the original data (with herbivory) and this simulation (no herbivory) estimated the influence of herbivory on average lifetime fitness for each subspecies at each site.

#### Simulation of fitness beyond the range edge with reduced herbivory

We were especially interested in the influence of herbivory on *xantiana* fitness beyond its range edge, but a complete absence of herbivory is unrealistic. Thus we used the same fitness predictions for eaten plants as above, but estimated mean fitness for both subspecies under the scenario where herbivory rates in the Beyond-Edge site were the same as in the Edge site (i.e., a reduction instead of complete removal of herbivory; details in Appendix S1). The lifetime fitness estimates for each subspecies in the Beyond-Edge site for both years were averaged over 100 simulations. Comparison of the predicted model means using the original data and this reduced herbivory simulation estimates the effect of increased herbivory rates outside the range limit on *xantiana* population persistence.

### To what extent does plant phenology mediate susceptibility to herbivory?

We predicted that differences in development rate between *parviflora* and *xantiana* contributed to the former’s escape from late season mammal herbivory at the Edge and Beyond-Edge sites (Table S1; Fig. S2), given observations that *parviflora* individuals are often dry and senescent when *xantiana* is still green and likely attractive to herbivores. Thus we tested whether plant phenology (as measured by flowering date) influenced a plant’s probability of late season herbivory, using data from the transplant experiment. Due to the very low survivorship and low incidence of herbivory in the dry year, we only analyzed these data for the wet year.

We were not interested in the date of flowering *per se*, but rather in using this as a proxy for a plant’s developmental speed. Thus, we predicted date of flowering for plants that died before flowering (from herbivory or other factors), enabling us to “recover” this missing phenological information and make more robust estimates of model parameters. Predictive models were evaluated using R^2^ statistics and diagnostic plots of predicted versus actual flowering dates (Appendix S1).

We tested the effect of date of flowering, with plant size and block as covariates, on a plant’s probability of fatal herbivory at each site using logistic regression with binomial error distribution and logit link. Because phenology is positively correlated with size in *C. xantiana* (Pearson’s *r* of log(size) and date of flowering = 0.47), we included size (here, the largest size a plant achieved) as a covariate in the models to isolate the effects of phenology. Plant size was calculated as the product of plant leaf number and average leaf length. Since we were interested in the relationship between phenology and late season herbivory only, these analyses were restricted to plants that survived early season herbivory (i.e., were alive at the March census); analyses including early season herbivory produced qualitatively similar results (Appendix S1). Since some plants for which we predicted flowering date died from factors other than herbivory (thereby precluding any later herbivory), these tests are somewhat conservative (i.e., some plants with predicted flowering dates were not eaten simply because they died before herbivores had the chance to eat them); in plots below we differentiate those plants that died from factors other than herbivory to assist in interpretation. We tested the significance of each term using Type II ANOVAs with likelihood ratio tests (car package; Fox & Weisberg 2011) and calculated model R^2^ (Nagelkerke’s pseudo R^2^) using the sjstats package (Lüdecke 2018). Conditional predicted probabilities were calculated and plotted with the visreg (Breheny & Burchett 2017) and ggplot2 (Wickham 2009) packages in R.

## Results

### Herbivore pressure increases at and beyond the range limit

In 2015, the probability of herbivory on translocated *xantiana* was low at the center of the range and increased sharply near the range limit, reaching levels above 0.75 outside the range limit (Fig. 2A). The pattern of herbivory was best fit with the logistic model including longitude (easting) as a linear term (BIC: 1324; N = 1278; Nagelkerke’s R^2^ = 0.49; Table S2). Overall, the odds of a plant being eaten increased 9% for every kilometer eastward, with the gradient in probability of herbivory becoming very steep near the range limit. For example, in the last census round, the probability of herbivory increased from 0.01 at the most central site to 0.13 10 km east of that site, but over the next 10 km eastward, increased to 0.7 approximately at the range limit. There was also a significant interaction of longitude with time (P < 0.001), with probability of herbivory increasing as the season progressed for range edge and beyond-range sites, but not at range center (Fig. 2A). Genotype (central vs. edge) had no effect on probability of herbivory (P = 0.5). Within *xantiana*’s range, herbivory on translocated stems generally matched that on natural plants, with rates at four of five sites differing by less than 5%; translocated stems experienced much more herbivory at one near-edge site but removing this site did not qualitatively affect the modeled gradient in probability of herbivory (see Appendix S1).

**Figure 2.**
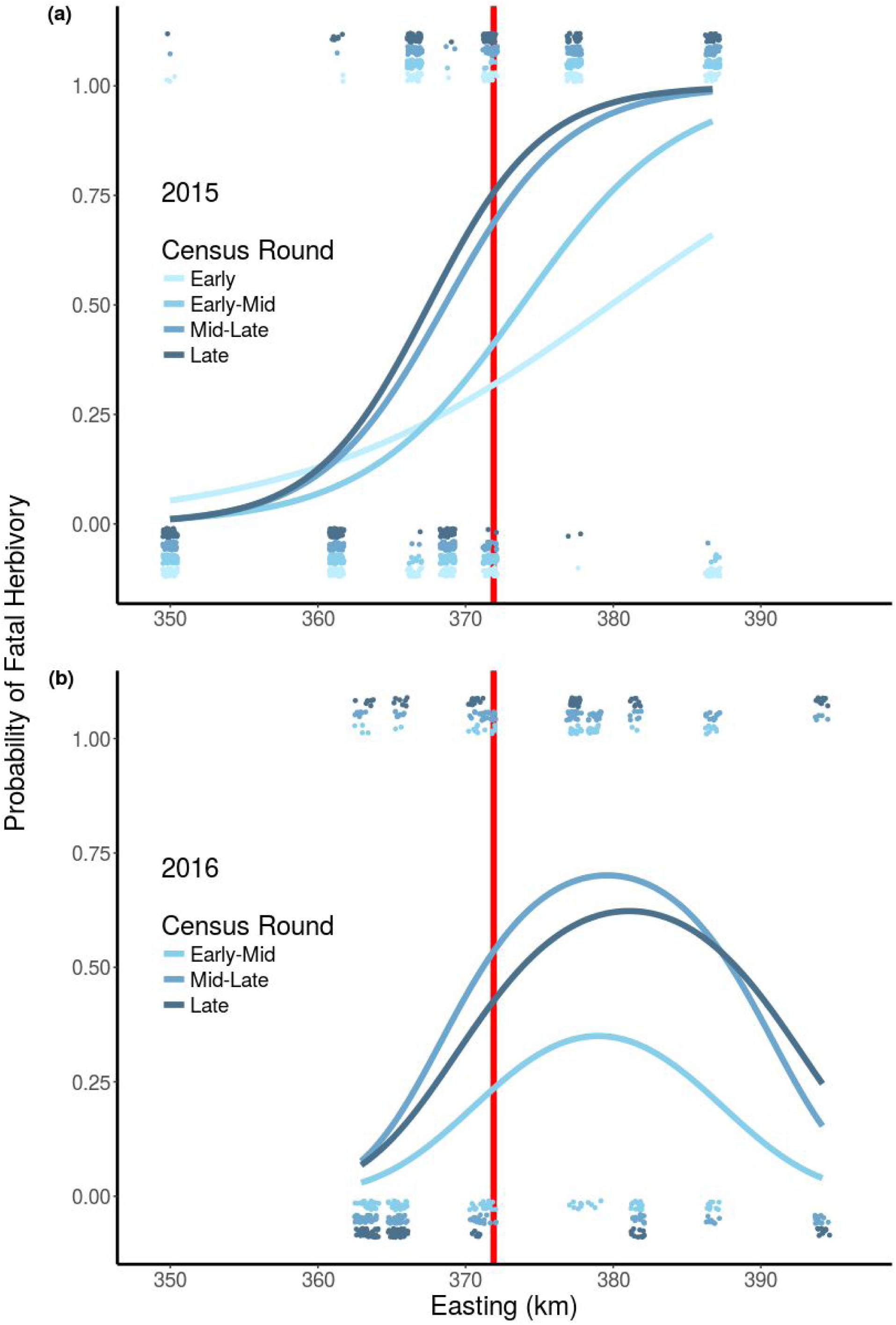
Probability of herbivory increases sharply near the range limit and over the growing season. (a) shows effects of Easting and Time on probability of herbivory as modeled by logistic regression for translocated *xantiana* stems in 2015, from generalized linear model of herbivory with easting, genotype, and transect as independent variables (Nagelkerke’s R^2^ = 0.49). Conditional effects of easting are shown for each census round, holding genotype (Central) and transect (one) fixed. Colors correspond to temporal replicates, which were installed approximately once per week 24 May - 19 June. Jittered points represent individual plants, which either did (located at top of plot) or did not (located at bottom of plot) experience herbivory. Vertical red line indicates location of *xantiana*’s eastern range limit. (b) shows effects of Easting and Time on probability of herbivory for translocated *xantiana* stems in 2016, from generalized linear model of herbivory with easting and transect as independent variables (Nagelkerke’s R^2^ = 0.33). Conditional effects of easting are shown for each census round, holding transect fixed at one. Colors correspond to temporal replicates, which were installed approximately once per week 5 - 25 June. Jittered points represent individual plants, which either did (located at top of plot) or did not (located at bottom of plot) experience herbivory. Vertical red line indicates location of *xantiana’s* eastern range limit.

In 2016, the pattern of herbivory was best fit with a logistic model including longitude as linear plus quadratic terms (BIC: 696; N = 561; Nagelkerke’s R^2^ = 0.33; Table S2). Probability of herbivory was low 10 km inside the range limit (~ 0.07), increased to a maximum of ~ 0.62 eight kilometers beyond the range limit, and decreased further east (Fig. 2B). Probability of herbivory also increased from the first census round to later rounds (P = 0.002), though there was no significant interaction with easting as in 2015.

### Herbivory threatens population persistence beyond the range limit

In the dry year of the transplant experiment, herbivory on both subspecies at all sites was very low (one to five percent of germinated plants eaten; Table S1). In the wet year, *xantiana* and *parviflora* suffered equal rates of herbivory (15%) at the Center site, but differing levels of herbivory at the Edge (*xantiana*: 34%; *parviflora* 8%) and Beyond-Edge *(xantiana:* 54%; *parviflora:* 19%) sites (Table S1).

When we simulated a scenario with no fatal herbivory, effects on fitness were observed in the wet year but not in the dry year, when plant survival and performance was low across all sites, and little herbivory was observed. In the wet year, removal of herbivory had the largest effect on lifetime fitness for *xantiana* in the Edge and Beyond-Edge sites, increasing lifetime fitness more than two and five fold, respectively, and *xantiana* fared slightly better at the Center site (Fig. 3; Table S3). Importantly, removal of herbivory beyond the range edge brought estimates of *xantiana* average lifetime fitness to 1, which suggests that populations could potentially replace themselves in the absence of herbivory. Removal of herbivory also increased estimates of *parviflora* fitness in these sites, though the effects were smaller (22% increase at the Edge and 120% increase Beyond-Edge).

**Figure 3.**
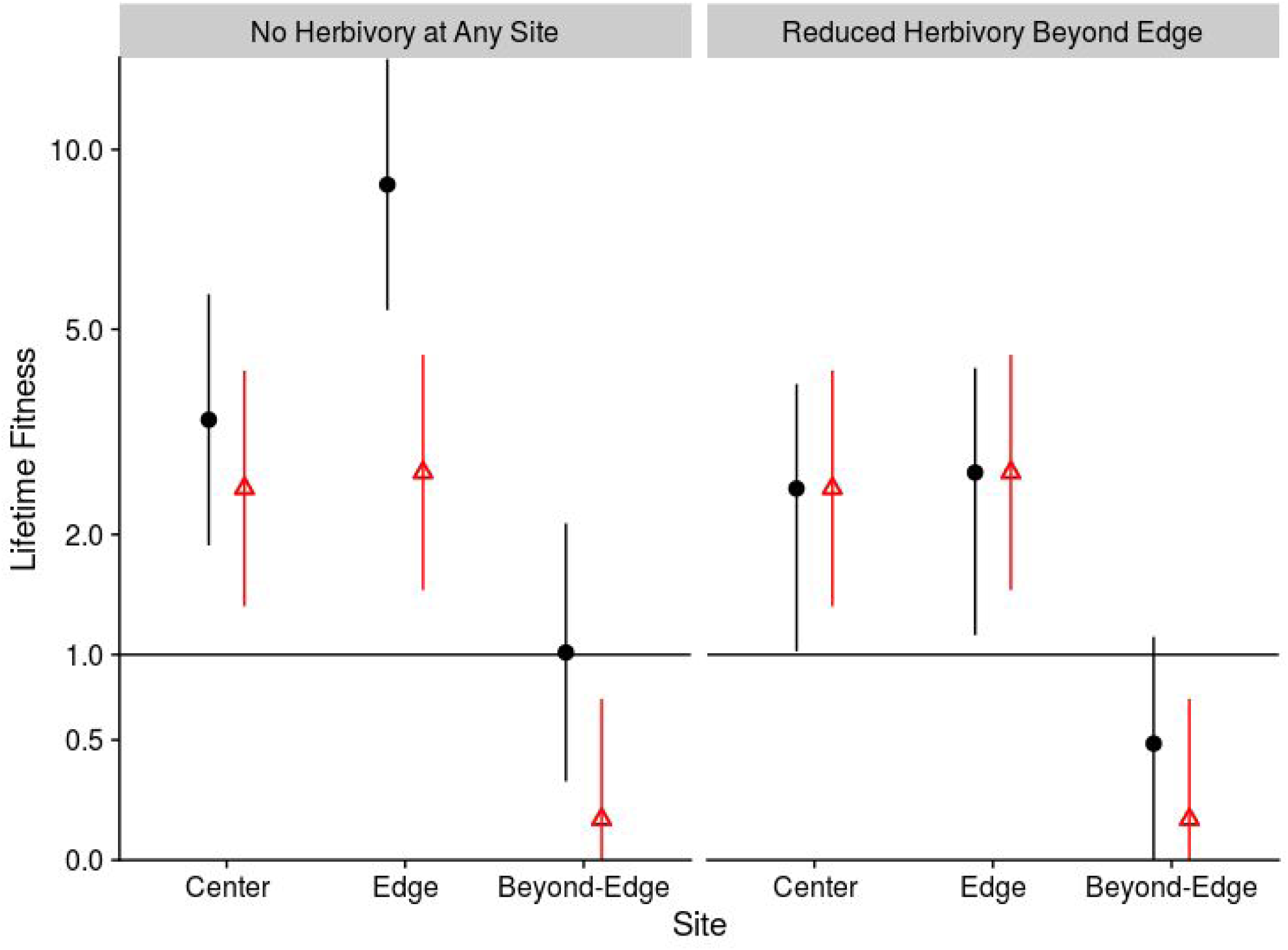
Removal and reduction of herbivory lead to large increases in *xantiana* fitness estimates at and beyond the range edge. Shown are lifetime fitness estimates for *xantiana* in each site in the Wet Year under two simulated scenarios: “No Herbivory at Any Site” simulation, where we predicted fitness values for all plants eaten during the field experiment as if they hadn’t been eaten; and “Reduced herbivory Beyond-Edge”, where we simulated lowered herbivory rates outside *xantiana*’s range limit (mimicking herbivory rates at the Edge site) but used the original data for Center and Edge sites. Black circles are lifetime fitness estimates from simulations; red triangles are lifetime fitness estimates using original field data (and thus are identical in both panels); point ranges show 95% confidence intervals. Note Y axis is on log scale. Results for *parviflora* are included in Appendix S1. Confidence intervals for points in the Reduced Herbivory simulation are averages from results of the 100 simulations; the CI’s for Center and Edge sites differ slightly from CI’s obtained from original model using field data due to slight differences in maximum likelihood estimates across the 100 simulated runs.

When we simulated a scenario where herbivory was reduced in the Beyond-Edge site to levels observed at the Edge site, *parviflora* and *xantiana* experienced increases in lifetime fitness estimates in the wet year but not in the dry year (Fig. 3; Table S3). In the wet year, average lifetime fitness for *parviflora* increased 50% to 3.54 and for *xantiana* increased 220% to 0.48.

### Phenology mediates late season herbivory

Phenology influenced probability of herbivory on *xantiana* and *parviflora* at all sites, but was especially important for explaining variation in herbivory at the Edge and Beyond-Edge sites (Fig. 4). For each day delay in flowering, a plant’s odds of herbivory in the range Center, Edge, and Beyond-Edge sites increased significantly by 2%, 5%, and 14%, respectively (Table S4). In the Edge and Beyond-Edge sites, larger plants were more likely to be eaten (P < 0.002); whereas in the Center site, smaller plants were more likely eaten (P < 0.001). Block was a significant factor in all sites (P < 0.001), indicating fine-scale spatial heterogeneity in herbivory. Differentiation in phenology between the two subspecies is illustrated in Figure 4, where the earlier phenology of *parviflora* is apparent. This difference in phenology translated to a marked difference in susceptibility to fatal herbivory between the subspecies at the Edge and Beyond-Edge sites.

**Figure 4.**
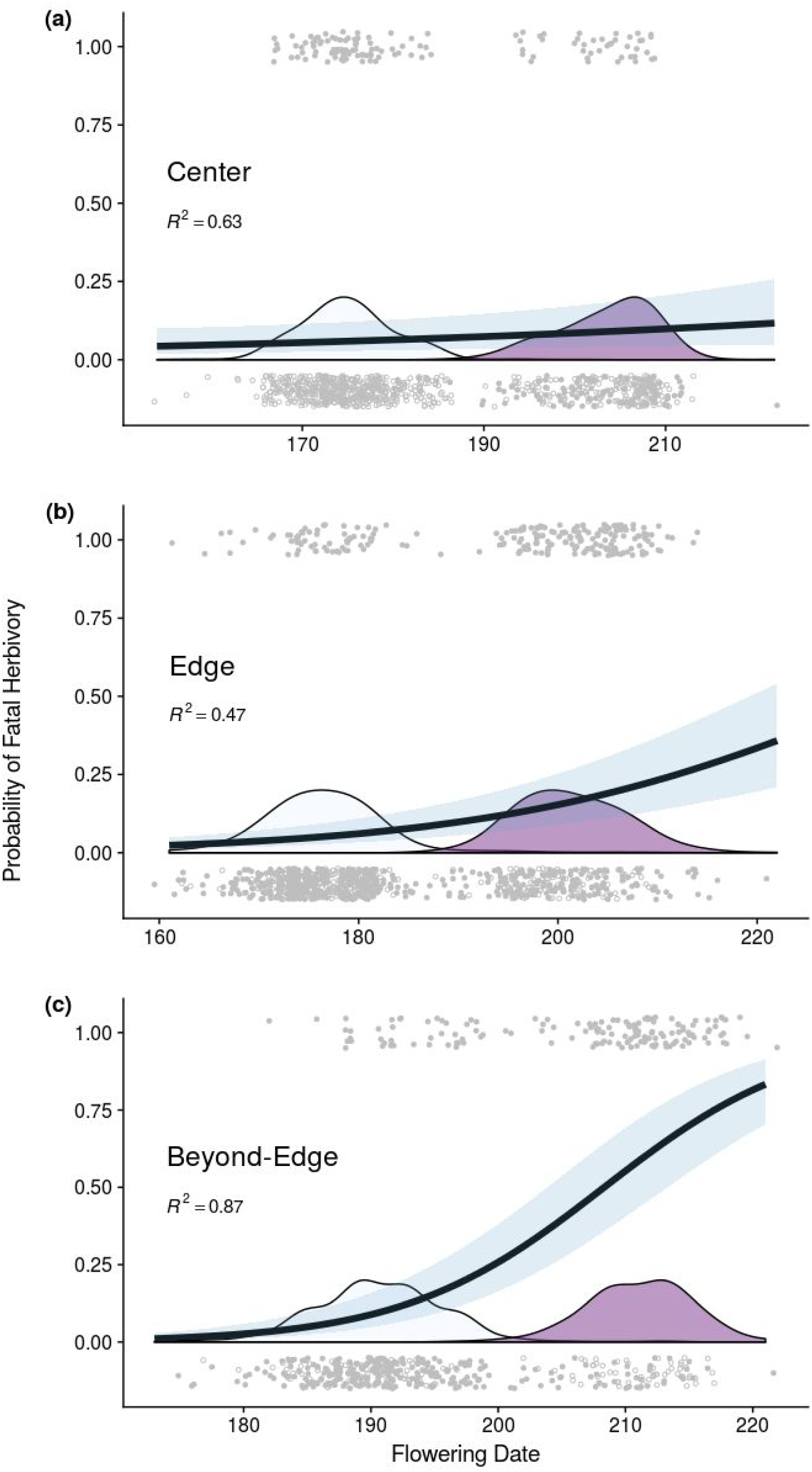
Phenology drives susceptibility to herbivory at and beyond *xantiana’s* range edge. Conditional effects of phenology (Flowering Date = days since 1 December) on probability of late season fatal herbivory as modeled by logistic regression, holding size and block constant, at Center (a), Edge (b), and Beyond-Edge (c) sites in the Wet Year, with 95% confidence bands. Note that X axes have different ranges due to overall site differences in phenology. Nagelkerke’s pseudo R^2^ is reported in each panel. Kernel density estimates (essentially, smoothed histograms) indicate distribution of flowering date for each subspecies (light blue = *parviflora*; purple = *xantiana*). Jittered points are individual plants that either did (at top of plot) or did not (at bottom of plot) experience herbivory. Open points indicate plants which died due to factors other than herbivory.

## Discussion

Recent reviews of transplant experiments support the idea that species’ geographic range limits often reflect niche limits (Hargreaves *et al.* 2014; Lee-Yaw *et al.* 2016). But given the demonstrated power of natural selection to produce adaptations to novel environments, what is preventing range expansion via sequential adaptation of marginal populations? The vast majority of work on geographic range limits has focused on gradients in abiotic variables, mainly temperature and precipitation. However, the field is increasingly calling for tests of how biotic interactions can modulate range boundaries, given experimental (e.g., Moeller *et al.* 2012; HilleRisLambers *et al.* 2013; Afkhami *et al.* 2014), theoretical (e.g., Hochberg & Ives 1999; Case & Taper 2000), and correlational (e.g., Araújo & Luoto 2007) evidence for the influence of species’ interactions on distributions at macroecological scales. Here we showed that an antagonistic biotic interaction, herbivory, has large effects on population mean lifetime fitness at and beyond the subspecies’ geographic range limit, and that probability of herbivory exhibits a steep gradient across the range of *C. xantiana.* We then showed that a specific plant trait, phenology, mediates probability of herbivory at and outside the range limit. Together, this set of results provides some of the strongest evidence to date that biotic interactions can play a pivotal role in determining the location of a geographic range limit.

Our simulations showed that at range center, removal of herbivory had minor effects on *xantiana* lifetime fitness, but at and beyond the range edge, simulating a complete absence of herbivory led to increases in estimates of *xantiana* lifetime fitness of two and five fold, respectively. For annual plants like *xantiana*, population mean lifetime fitness approximates population growth rate (λ). Interestingly, this meant that the estimate of *xantiana* population growth at the range edge was more than double that at range center in the absence of herbivory. This highlights how a biotic interaction can influence population demography at a species’ range edge, and potentially emigration and colonization outside the range limit.

We also showed that without herbivory, populations beyond the range edge could potentially replace themselves, providing some of the strongest evidence to date that a biotic interaction can limit a taxon’s geographic range. When we instead reduced herbivory outside the range, *xantiana* mean lifetime fitness increased 220% relative to field data in the wet year, to *λ* = 0.48. Though this is still below levels needed for population replacement, adaptive evolution beyond the range margin could raise population mean fitness above replacement, given adequate heritable variation in ecologically important traits. There is evidence of substantial genetic variance for fitness in *xantiana* planted beyond its range limit (M. Geber, unpubl. data), and if this allowed population mean fitness to evolve then populations could theoretically “escape” extirpation as *λ* values climb toward 1 (Fisher 1930; Shaw & Shaw 2014).

Our stem translocation experiments showed that herbivory exhibits a particularly steep gradient across and beyond *xantiana*’s range, with a sharp increase in probability of herbivory near the eastern range margin. For example, during the last stem census in 2015, plants at the center of the range had less than a five percent chance of fatal herbivory, while only eight kilometers outside its range limit, probability of herbivory on *xantiana* went up more than fifteen-fold. By comparison, mean growing season precipitation across that same expanse decreases about twofold (Eckhart *et al.* 2010), and pollinator abundance decreases roughly five-fold (Moeller 2006).

These phenomena speak to the proximate, ecological causes of *xantiana’s* range limit, but the ultimate cause of a range limited by adaptation is genetic limits on trait evolution. We rarely know which traits would need to evolve to allow range expansion (but see Hoffmann *et al.* 2003; Griffith & Watson 2006; Angert *et al.* 2008; Colautti *et al.* 2010). In this study, we were able to use differentiated sister taxa to ask how a specific trait, phenology, influenced probability of herbivory at multiple sites. While phenology had little effect at range center, the difference in phenology between the two subspecies beyond the range limit drove strong differences in susceptibility to fatal herbivory. Phenology has been shown to be a key range-limiting trait in other plant species, though usually in the context of abiotic latitudinal range limits (Griffith & Watson 2006; Colautti *et al.* 2010). For *xantiana*, it seems phenology would have to evolve to enable eastward range expansion, and thus the question becomes, why has it not?

Recent theoretical work by Polechová and Barton (2015) showed that in models including genetic drift, a range margin can form via two (non-mutually-exclusive) mechanisms: a steepening (i.e., non-linear) environmental gradient driving increasing maladaptation, or a decrease in carrying capacity across space leading to an increased influence of drift on population genetic variance. Both of these factors could be at play for *xantiana.* In these models, a steepening environmental gradient creates a sharp range margin near the environmental “inflection point” due to drift eroding genetic variance needed to adapt to a quickly changing trait optimum. The result is that population trait means closely track trait optima along most of the environmental gradient, but fail to do so when this gradient suddenly steepens, like the gradient in probability of herbivory does near *xantiana’s* range limit. Estimating “optimal” flowering dates for the three transplant sites (Appendix S1) and comparing these optima to the actual mean flowering date for *xantiana* shows that *xantiana* is very far from the phenological optimum (18 days later) outside its range, but is within 4 days of the estimated optima at Center and Edge sites (Fig. S9). Increased herbivore pressure could also impose an extrinsic limit on *xantiana*’s carrying capacity outside its range edge, depressing population sizes so as to make any populations able to colonize outside the range limit more susceptible to drift eroding potentially adaptive genetic variance. The concordance of observed patterns of environmental variation and *xantiana*’s distribution with model predictions provide empirical support for recent range limit models (Polechová & Barton 2015).

## Why does herbivory vary across space?

Geographic variation in herbivory across *xantiana*’s range can be explained by two phenomena. First, the herbivore community likely changes across *xantiana*’s range. The two primary lagomorph herbivores in this study area are the Desert cottontail (*Sylvilagus audubonii*) and the Black-tailed jackrabbit (*Lepus californicus*). Motion-triggered cameras installed at some sites outside the range limit in 2015 and 2016 captured both lagomorph species (and to a lesser extent, small rodents) preying on translocated stems (Fig. S1C). However, we have yet to observe *L. californicus* in more central *xantiana* habitat, where we often see *S. audubonii.* Habitat descriptions support these observations, reporting that *L. californicus* is more common in arid, open scrub land typical of sites outside *xantiana*’s eastern range boundary (Arias-Del Razo *et al.* 2012). If there is increased herbivore pressure outside *xantiana*’s range limit due to an additional lagomorph species, this could translate into higher herbivory rates on *xantiana* planted outside its range limit.

A second, non-mutually-exclusive hypothesis is based on decreases in primary productivity, especially of forbs, across the west-to-east gradient. With more forage available at *xantiana’*s range center, this may dilute herbivore pressure on *xantiana*, whereas in the more arid east, *xantiana* may be increasingly attractive to herbivores due to limited forage and its late completion of development compared to co-occurring forbs. Field observations suggest this pattern arises because *parviflora* is less palatable forage by the peak of late season herbivory, whereas *xantiana* is still green and flowering. For example, during transplant experiments, *xantiana* was often the only herbaceous vegetation still green by early June, when surrounding ephemerals had already senesced.

## Generality of a generalist predator enforcing range limits

Given the strong effects of herbivory on individual plant fitness, population growth, and local distributions (Louda 1982; Quinn 1986), it is surprising that no studies have examined herbivory’s role in modulating plant species’ geographic ranges (Maron & Crone 2006; but see Bruelheide & Scheidel 1999). Case et al. (2005) pointed out that, theoretically, polyphagous predators can easily enforce geographic range limits of prey species, especially when two prey species are differentially susceptible to predation over a spatial gradient. This is the pattern we see in *C. xantiana*, but should we expect that generalist herbivores often regulate geographic distributions of plant species? Rapid phenology is commonly observed in arid systems, and this has long been presumed to be due to selection to escape the late season drought and unpredictable hydric environments of arid areas (Aronson *et al.* 1992; Thuiller *et al.* 2004; Levin 2006; Volis 2007). “Phenological escape” from insect herbivory has been shown for multiple plant taxa (Pilson 2000; Krimmel & Pearse 2016; Mlynarek *et al.* 2017), but mammalian herbivore control on plant phenology and distributions in arid environments remains relatively unexplored.

We often focus on climatic control of geographic range limits, but given the relatively constant, gradual nature of most climatic gradients, and predictions from recent theoretical models, we might expect that ranges limited by adaptation are not often set by these sorts of environmental gradients alone. Combining multiple lines of evidence to link environmental variation, traits, and fitness, our study demonstrates how biotic interactions can generate adaptive hurdles for important traits and contribute to the formation of species’ range limits.

## Acknowledgements

We thank C. McGuire, M. Meixner, M. Patel, J. Runions, E. Twieg, J. Hanson, S. Southard, J. Fausto, S.Travers, and C. Noss for assistance in the lab and field during the transplant experiment, and L. Bolin for assistance during the stem-translocation experiment. The manuscript was greatly improved with comments from the Moeller Lab, and we thank J. Polechová for correspondence relative to recent theoretical models of range limits.

